## Supplementary figures 1-9 and table 1 for "Reversible DNA condensation drives natural transformation"

This PDF file includes:

Figs. S1 to S9, Table S1

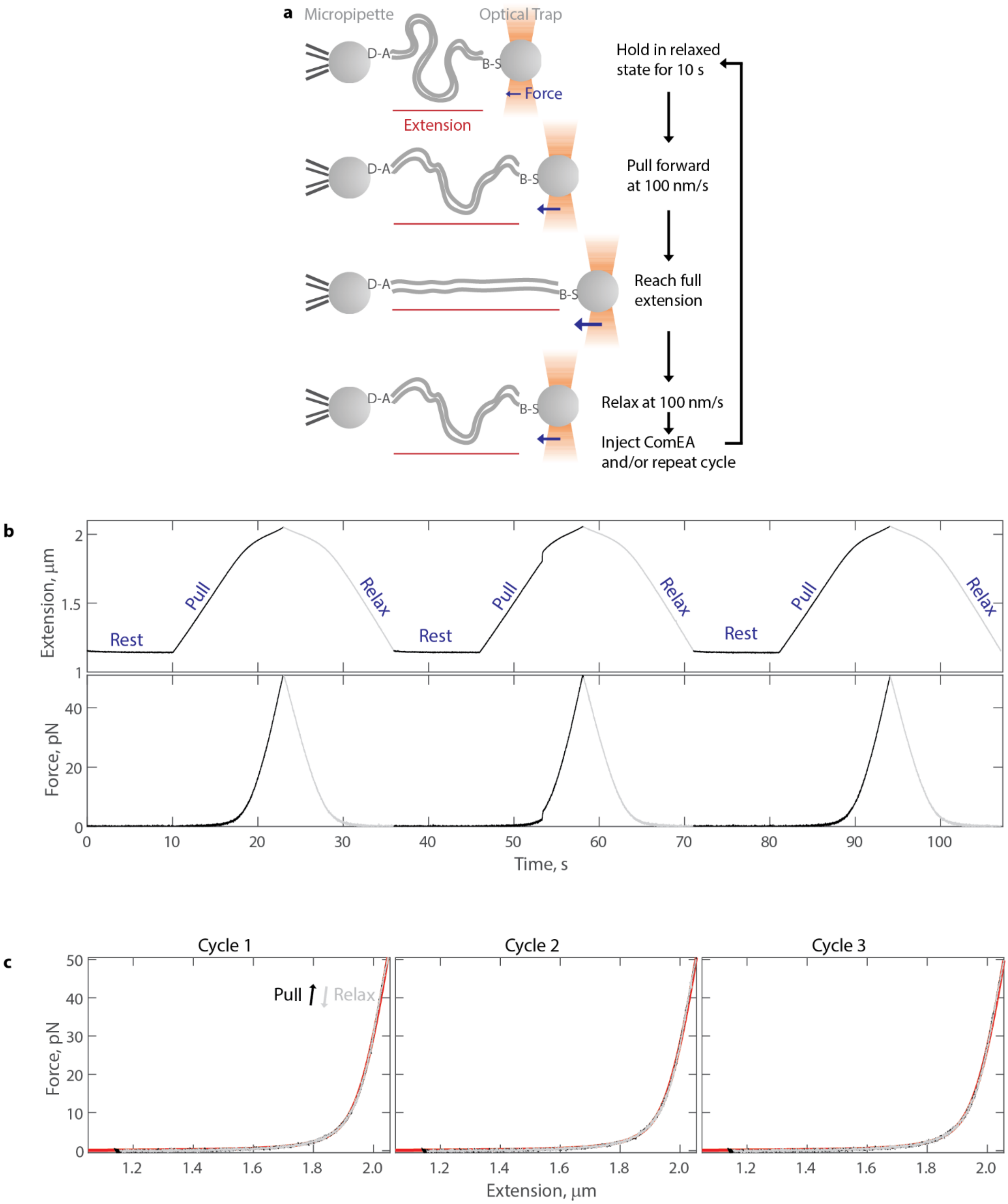


**Figure S1. Detailed force extension cycle**

(a) Diagram of optical tweezers force-extension assay. The DNA is connected to a DBCO (D) bead via a 5' azide (A) and to a streptavidin (S) bead via a 5' biotin.

(b) Data showing the extension of 6 kbp DNA over time following the protocol in panel a. Shown below is the resulting force over time.

(c) Data from panel b replotted as force extension curves. Rest and pull phases in black, relax phase in gray. The red line shows a fit to the extensible worm-like chain equation.

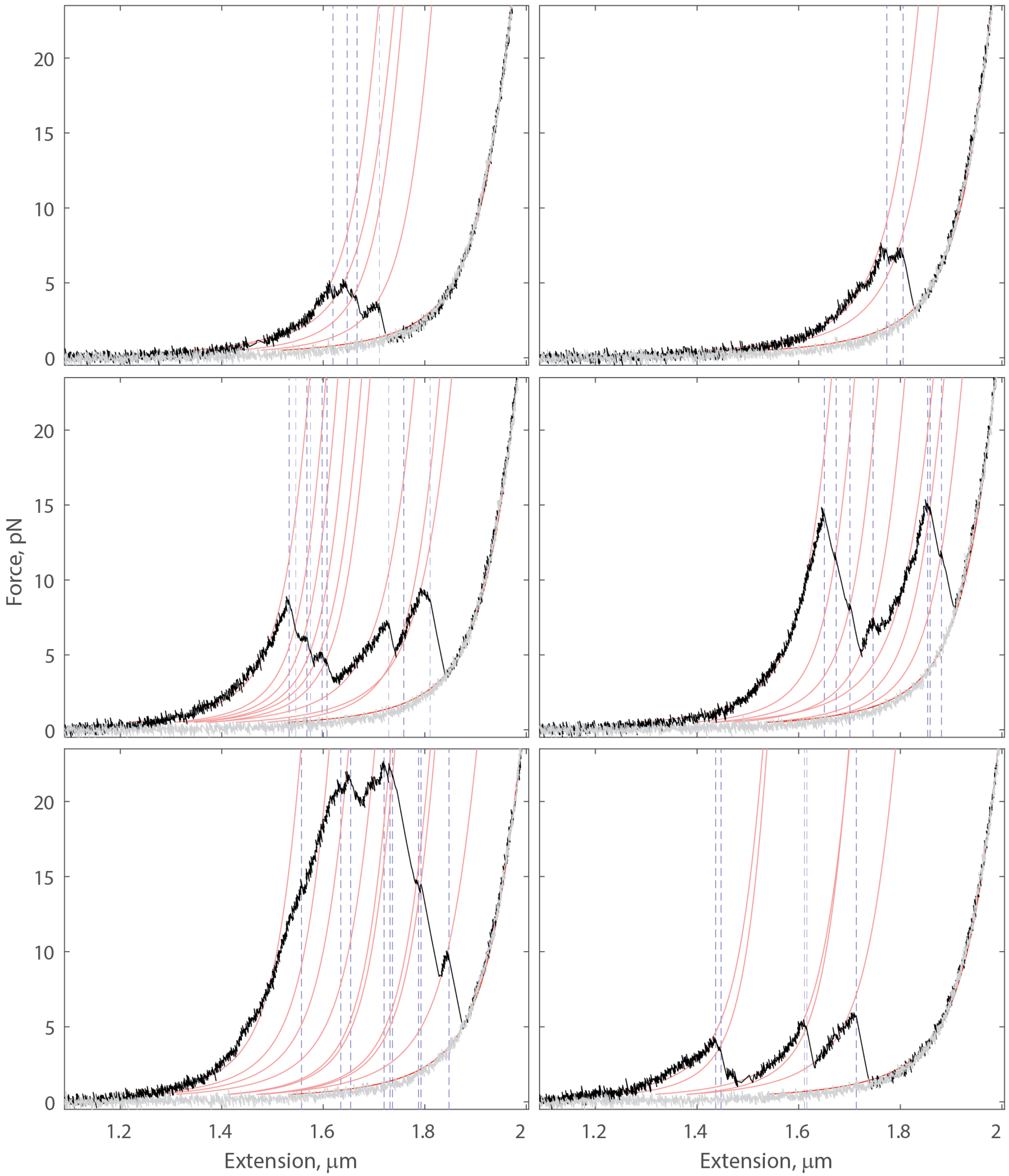


**Figure S2. Additional examples of force extension curves on 6 kbp dsDNA with 200 nM ComEA**
Six successive cycles with rest and pulling phases in black and relaxing phases in gray. Blue dotted lines show positions where oligomer breakage events were detected. Red curves show attempts to fit the data in between sets of blue dotted lines to the extensible worm-like chain model.

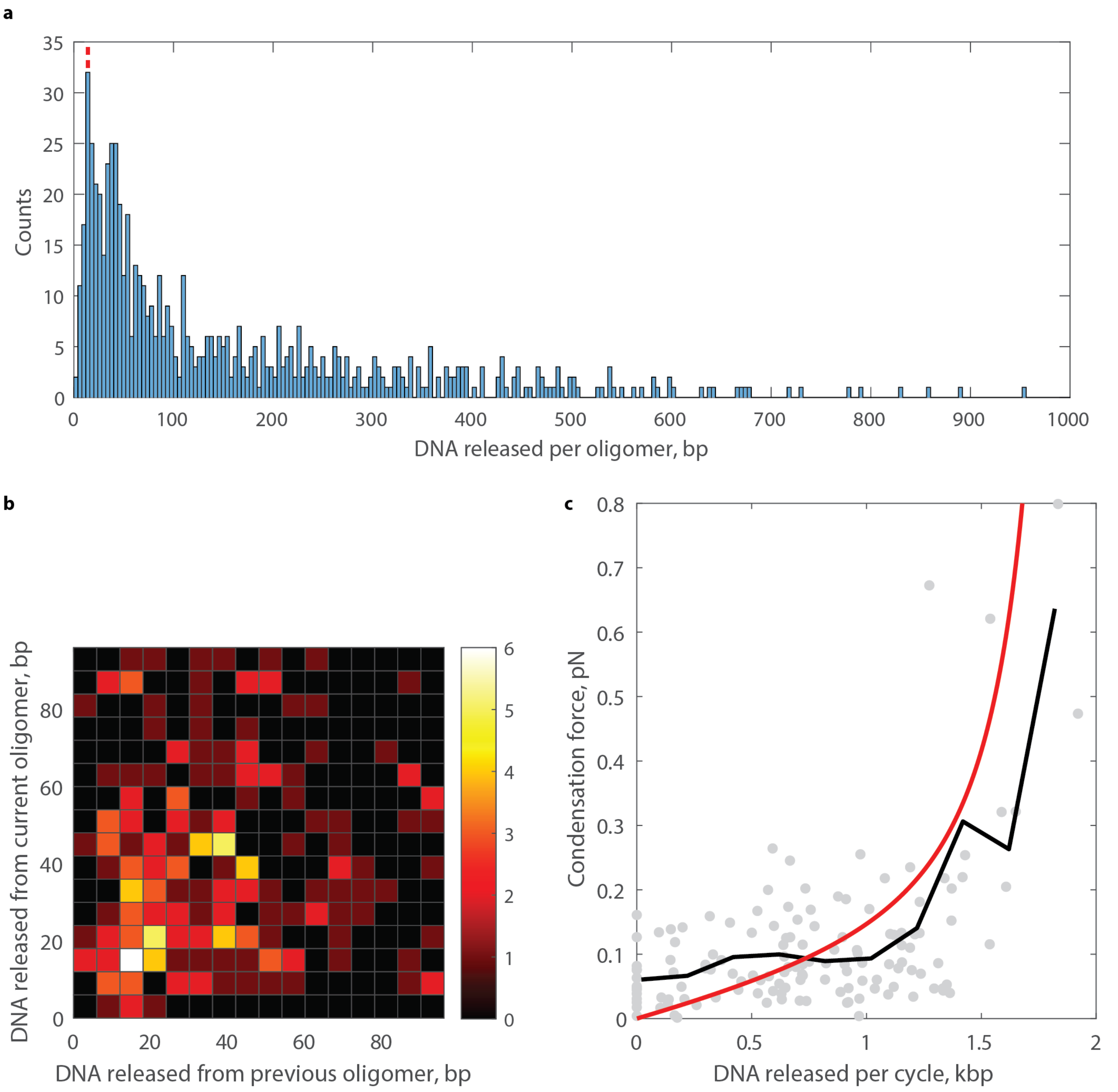


**Figure S3. Analysis of DNA released and force generated for force extension curves on 6 kbp dsDNA with 200 nM ComEA**

(a) Histogram of the amount of DNA released by each individual observed oligomer breaking (N=671 events). The red dotted line indicates 14 bp, equal to two ComEA DNA binding footprints.

(b) Histogram of the DNA released by a given oligomer based on how much DNA was released by the previous oligomer. This analysis ignores the first oligomer broken in each cycle. The highest counts are observed in the bin centered at 14 bp by 14 bp.

(c) Plot of the measured condensation force during the rest phase of each cycle versus the measured DNA released per cycle of the pulling phase of that cycle (N=144). These independent measurements scale accordingly. The black line shows the average condensation force in 200 bp bins along the X-axis. The red line shows the theoretical extensible worm-like chain equation for 6 kbp DNA.

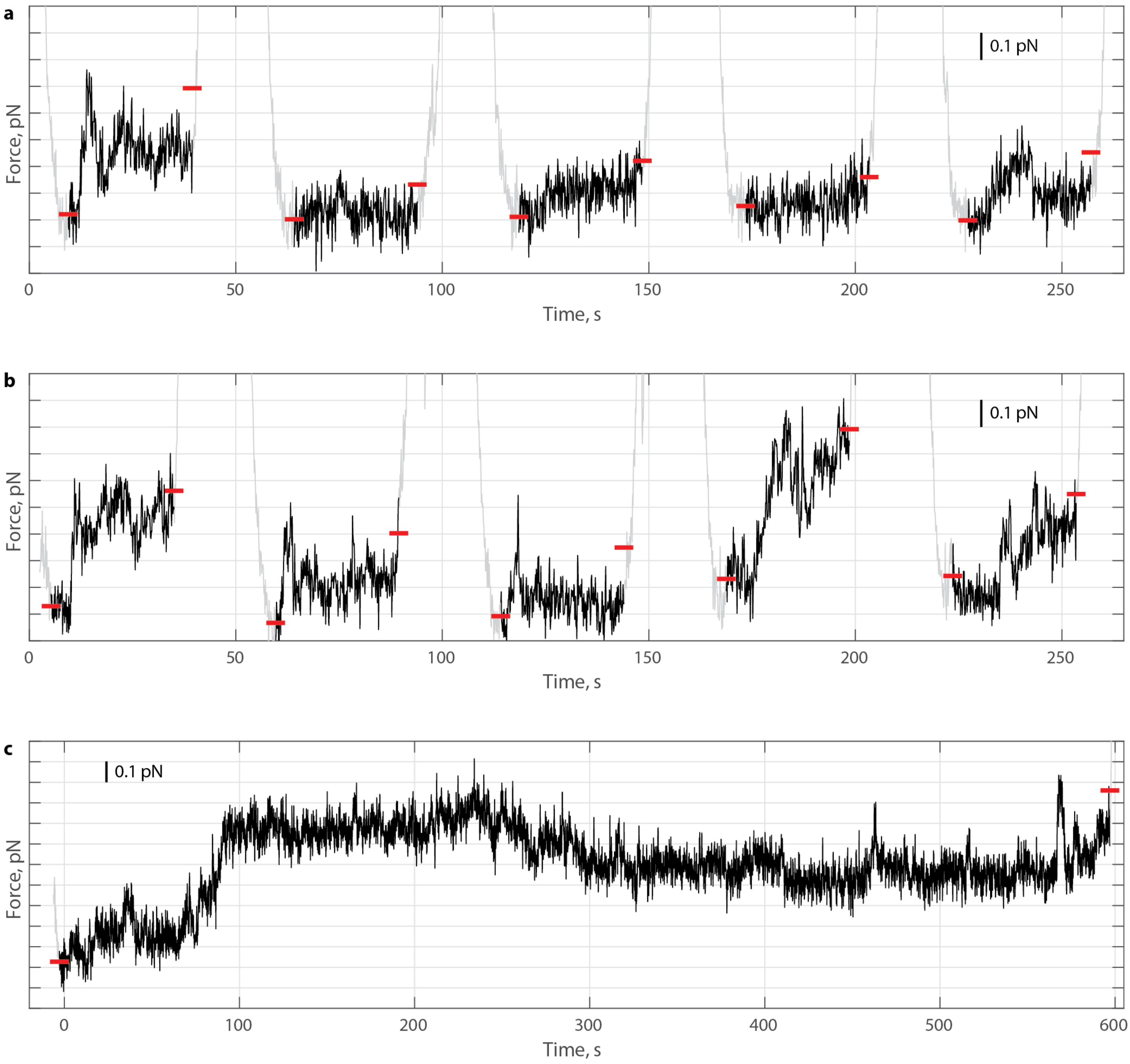


**Figure S4. Additional examples of force production during force extension curves on 6 kbp dsDNA with 200 nM ComEA with elongated rest phases**

(a) Example force versus time data during the 30-second rest phase of seven successive force-extension cycles after injecting 200 nM ComEA. The rest phases are colored black, while the pulling and relaxation phases are colored gray. Horizontal red dashes denote the average force during the first second of rest (left side) and the highest force held for one second during the rest phase (right side).

(b) Additional example, same conditions as panel a.

(c) Example data showing a force extension cycle with a 600-second rest phase.

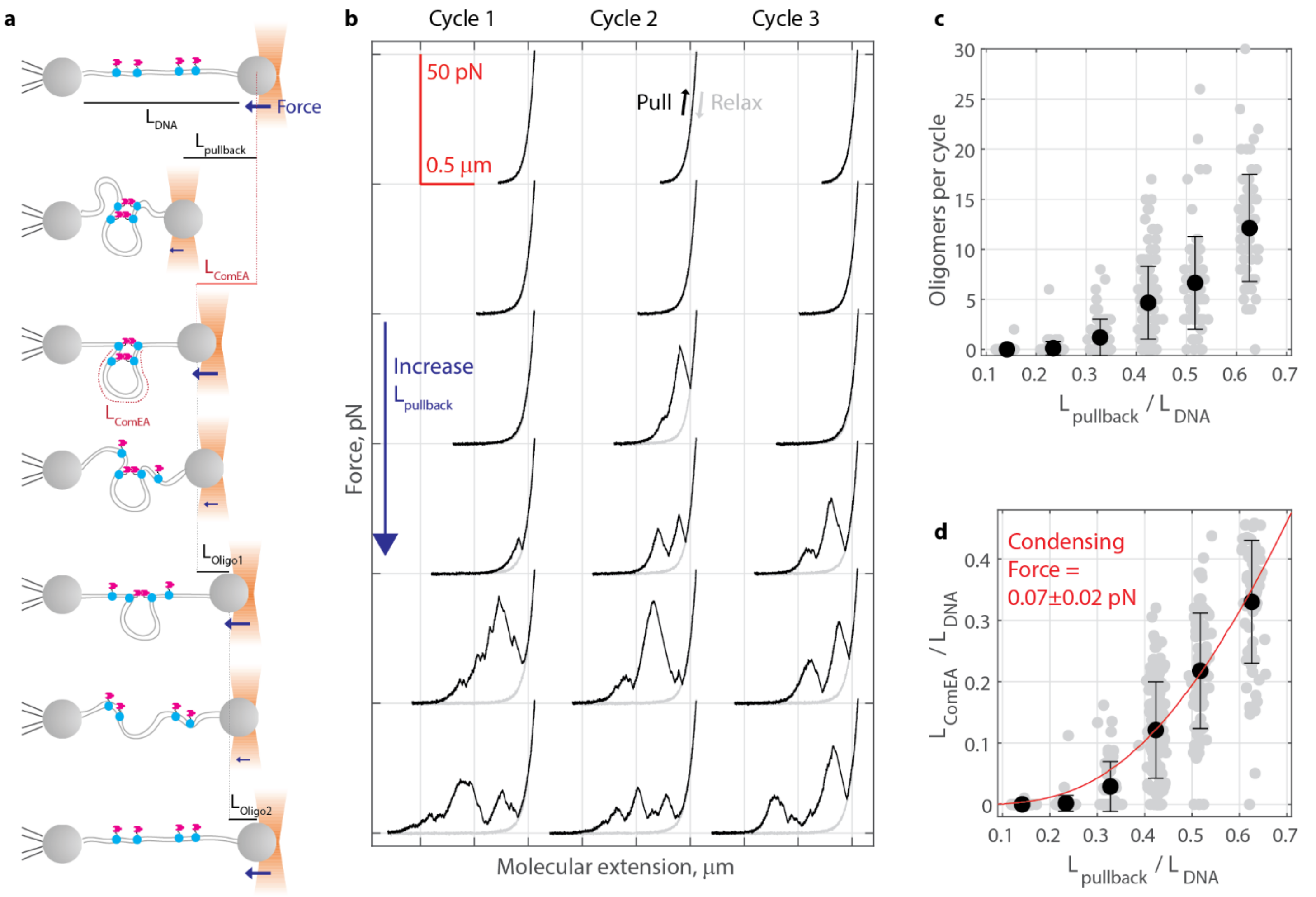


**Figure S5. Varying the pullback distance leads to dramatically different DNA condensation results**

(a) Diagram showing experimental design and definition of events. A strand of DNA with contour length $L_{DNA}$ is fully extended, then pulled back some distance $L_{pullback}$. The trap is help at that position for 10 seconds, then pulled forward again. The total amount of DNA looped off (condensed) by ComEA, $L_{ComEA}$, is determined via the extensible worm-like chain equation using the difference in extension at the point of the first oligomer breaking versus the extension on naked DNA at an equivalent force. The subsequent amount of DNA released by each oligomer is calculated similarly.

(b) Example data from a single strand of DNA in the presence of 200 nM ComEA. A variety of $L_{pullback}$ values were measured. Differences between the pulling phase (black) and relaxation phase (gray) are indicative of ComEA activity; each sudden, diagonal drop in force indicates the release of DNA due to oligomers breaking.

(c) The number of oligomers observed per cycle at six different $L_{pullback}$ values tested. Individual measurements are shown as a gray dot (slightly offset in X for clarity); mean values ± standard deviation (N = 113, 82, 76, 63, 142, 82, 72 cycles from left to right, respectively) are shown in black.

(d) The total amount of DNA released by ComEA per cycle as a function of $L_{pullback}$; each value normalized to the total DNA length (% pulled back versus % captured by ComEA). Individual measurements are shown as a gray dot (slightly offset in X for clarity); mean values ± standard deviation (N = 113, 82, 76, 63, 142, 82, 72 cycles from left to right, respectively) are shown in black. The data were fitted using the extensible worm-like chain equation, assuming ComEA only changes the DNA contour length and that its condensation activity does not substantially move the trapped bead (red curve).

Fit performed:

$$\begin{cases} x(F_1) = L_{DNA} - L_{pullback} = L_{DNA}\left(1 - 0.5\left(\frac{k_B T}{F_1 P}\right)^{0.5} + \frac{F_1}{S}\right) \\ x(F_2) = L_{DNA} - L_{pullback} = (L_{DNA} - L_{ComEA})\left(1 - 0.5\left(\frac{k_B T}{F_2 P}\right)^{0.5} + \frac{F_2}{S}\right) \\ Condensation\ force = F_2 - F_1 \end{cases}$$

assuming P=45 nm, S=1200 nm., $L_{DNA}$=6000 bp (2040 nm), and T=25̊ C. Fit weighted by inverse standard error of the mean. The fitted condensation force was 0.07 ± 0.02 pN (fit ± 95% confidence intervals determined by bootstrapping).

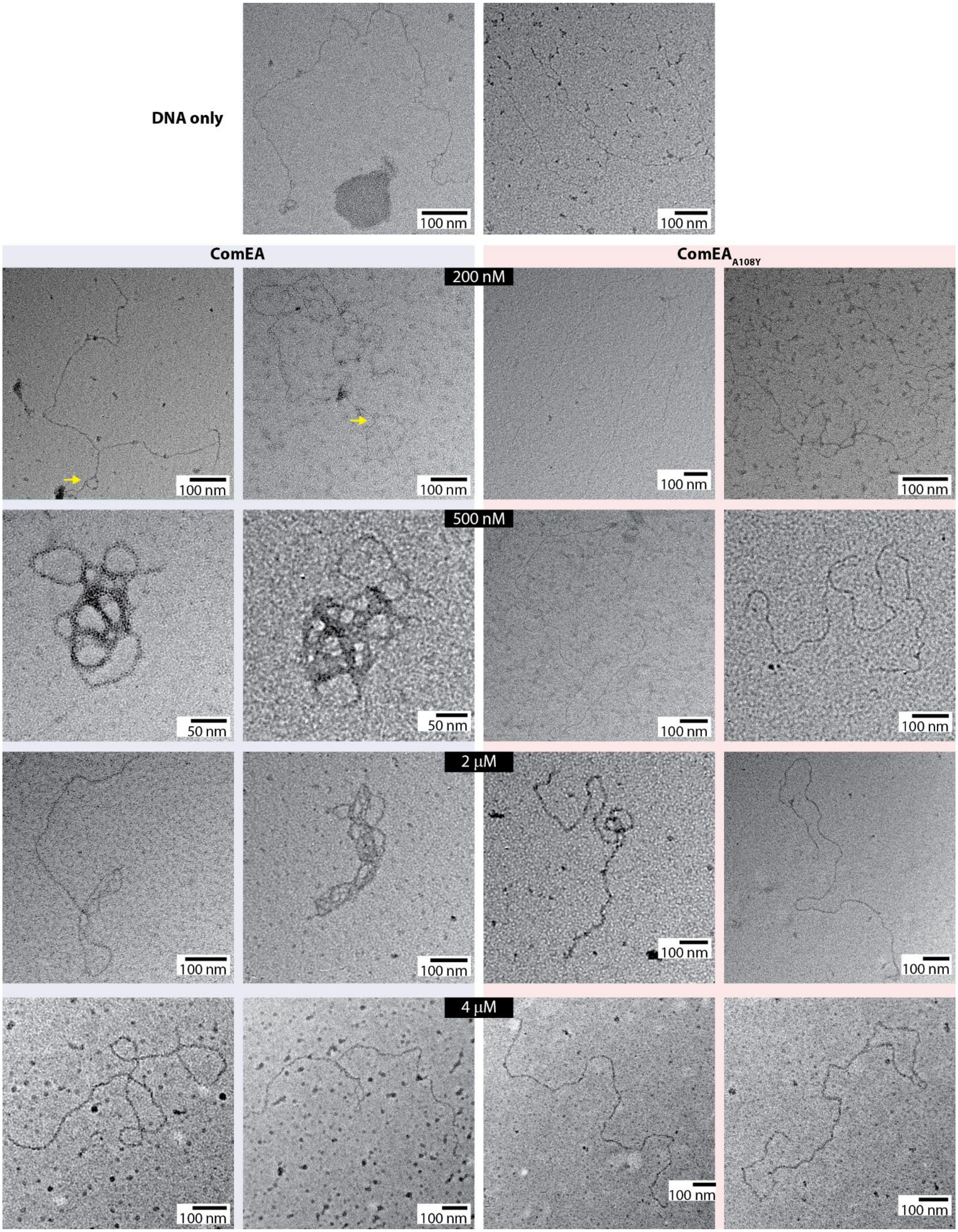


**Figure S6. Additional example negative stain electron micrographs of DNA condensed by ComEA**
Example negative stain electron microscopy images of linearized 5.495 kbp dsDNA with the labeled amount of ComEA or ComEA$_{A108Y}$. Yellow arrows denote loop structures with extended necks.

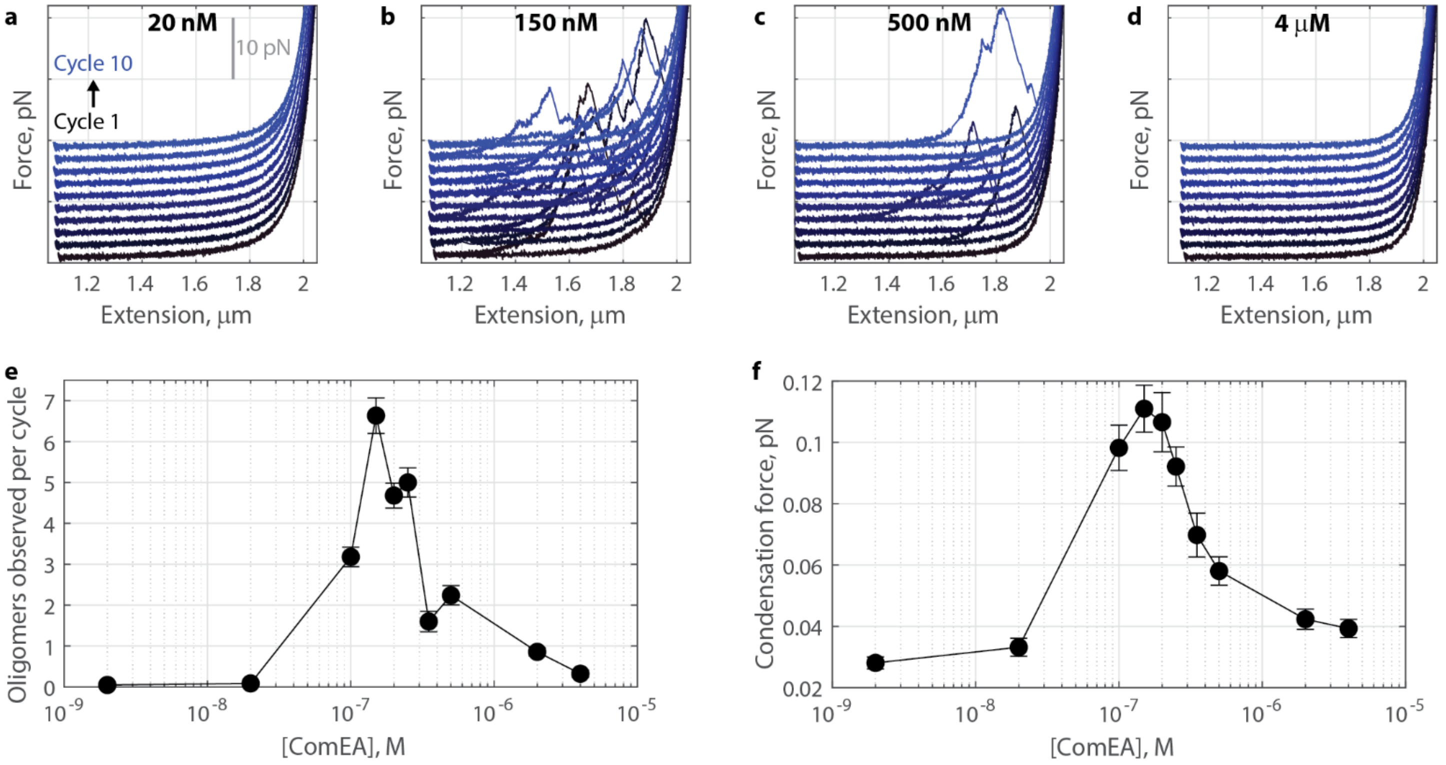


**Figure S7. ComEA DNA-bridging oligomer formation and force generation are concentration-dependent.**

(a) Ten successive force-extension curves (including 10 s rest phase) of DNA after injecting 20 nM ComEA. Each cycle is slightly offset on the force axis and colored for clarity.

(b-d) Example force extension curves, as in panel a, after injecting 150 nM, 500 nM, or 4 μM ComEA.

(e) The number of oligomers observed per cycle at various ComEA concentrations. Data plotted as mean ± standard error of the mean for n=106, 105, 100, 96, 144, 94, 75, 100, 98, and 90 cycles off N=5, 3, 10, 4, 14, 8, 7, 6, 4, and 7 molecules of DNA for 2 nM, 20 nM, 100 nM, 150 nM, 200 nM, 250 nM, 350 nM, 500 nM, 2 μM, and 4 μM ComEA, respectively.

(f) The condensation force generated during the rest phase for each ComEA concentration. Data plotted as mean ± standard error of the mean, same n and N as panel e.

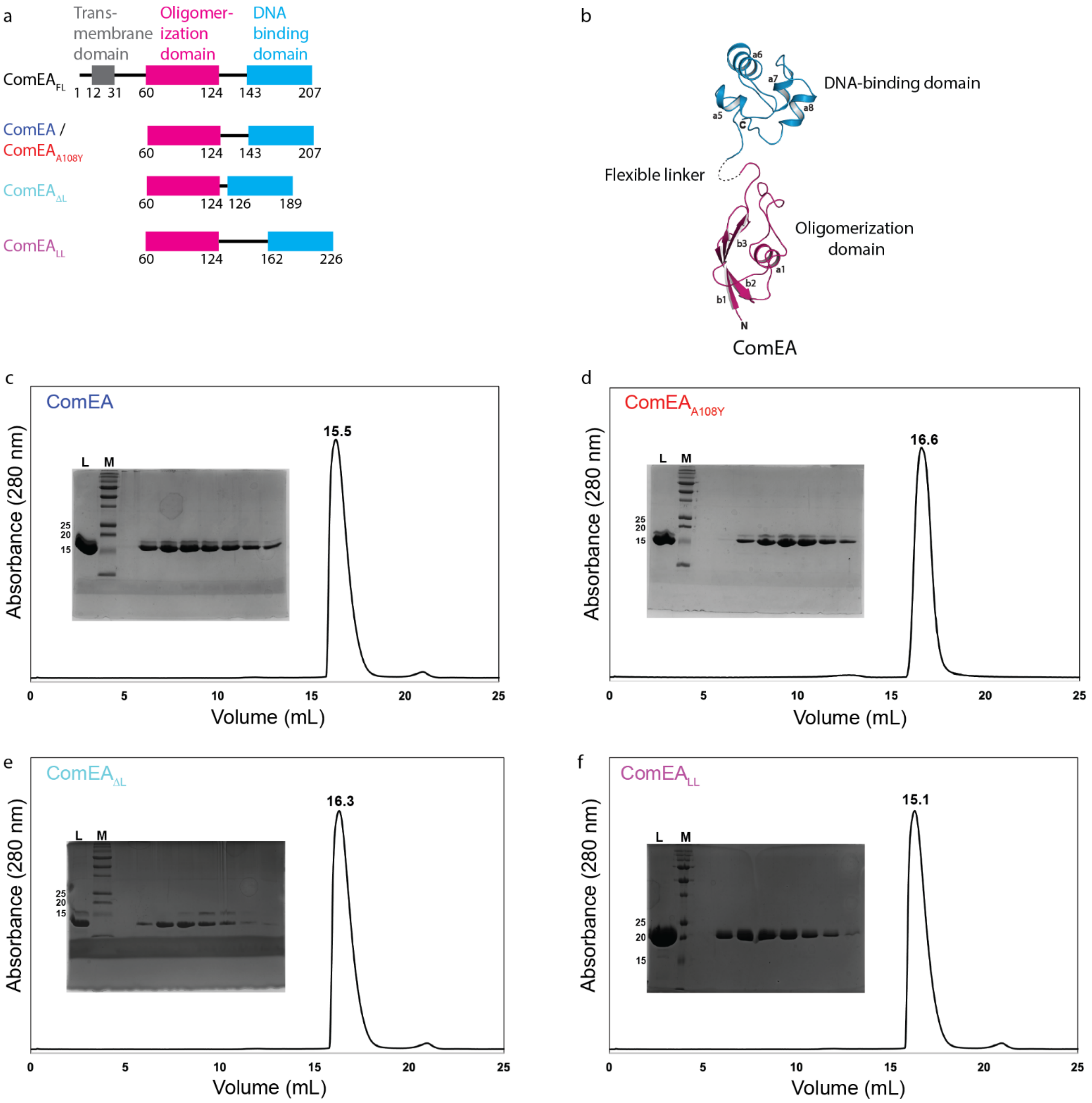


**Figure S8. Size exclusion chromatography analysis of wild-type ComEA, ComEA-A108Y, and ComEA linker deletion and insertion mutants.**

(a) Domain diagrams of ComEA and its mutants.

(b) Previously-solved crystal structure of ComEA (PDB 8DSS). Amino acids 62-124 are colored magenta, and 143-207 are colored cyan. The linker region (125-142) lacked density in this structure.

(c-f) SEC profiles of wild-type (WT) ComEA and its mutants were using a Superdex 200 increase 10/300 GL column. c-f show the SEC profiles for WT ComEA, $ComEA_{A108Y}$, $ComEA_{\Delta L}$, and $ComEA_{LL}$, respectively. The inset images in each panel show SDS-PAGE analysis of fractions collected across the peaks. Lanes include the loaded sample (L) and molecular weight markers (M). The 15, 20, and 25 kDa molecular weight markers are indicated. Experiments were repeated at least twice with similar results.

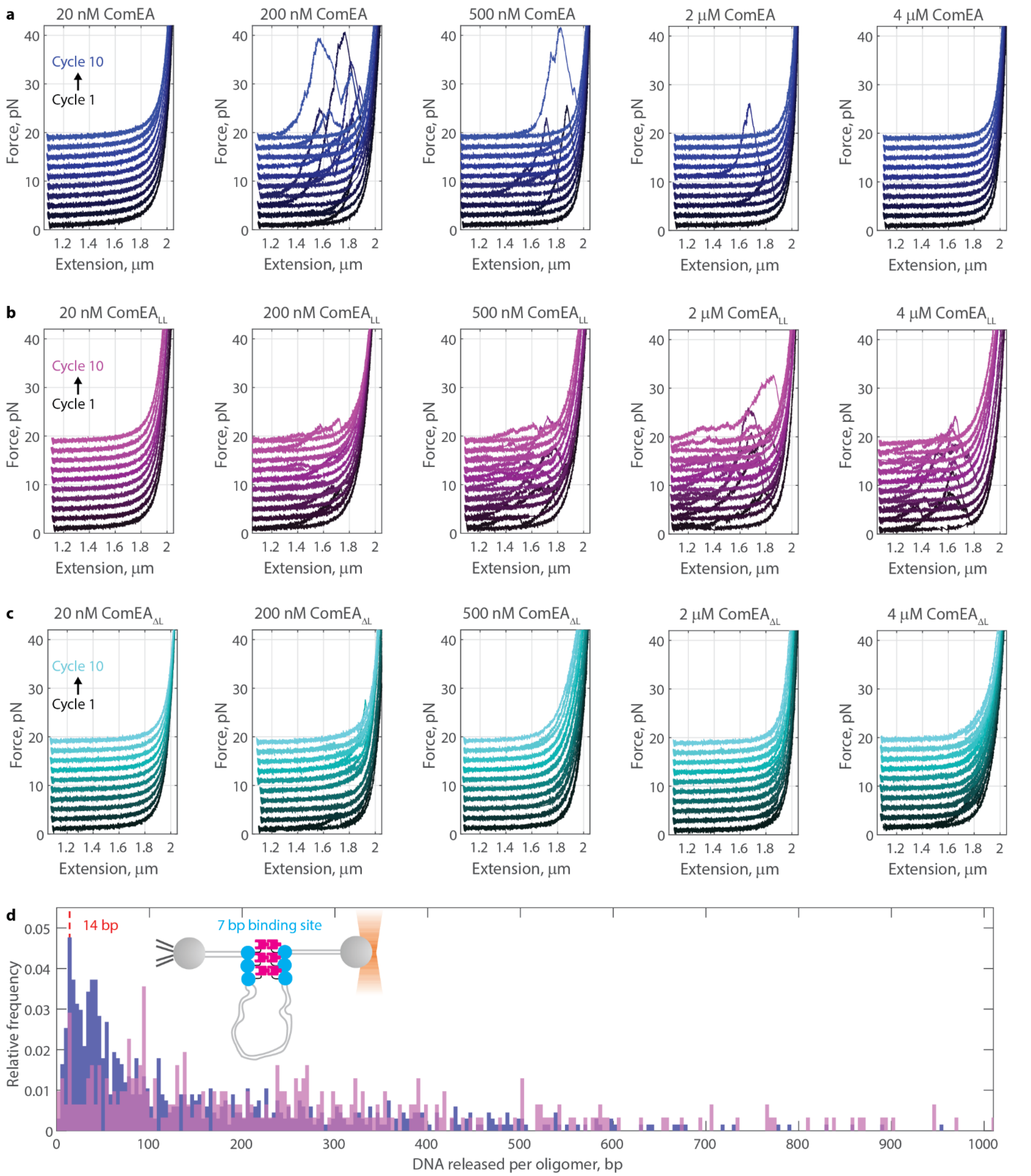


**Figure S9. Optical tweezers force-extension data for ComEA linker mutants**

(a-c) Example force extension traces for ComEA and linker mutants. Ten successive cycles are slightly offset on the force axis for clarity. Each plot is labeled with the concentration of ComEA added.

(d) Histogram of DNA released per oligomer for 200 nM ComEA (blue; n=671) and 200 nM $ComEA_{LL}$ (magenta, n=309). Both have a prominent peak at 14 bp. One interpretation is that ComEA molecules, which have a 7 bp binding footprint, line up at the base of loops (diagram inset).

**Table S1. Oligonucleotides used in this study**

| | |
|---|---|
| $EA_{Gs\Delta L}$ Fwd | 5′- GGGATGCAAGTGGCCATCAATACGGC -3′ |
| $EA_{Gs\Delta L}$ Rev | 5′- CTCGCCTTTTTTCGGCACGTAAATCAT -3′ |
| $EA_{GsLL}$ Fwd 1 | 5′- CGATGGTGCACCTAAGAACATGCAAGTGGCCATCAATACGGCGACGG -3′ |
| $EA_{GsLL}$ Fwd 2 | 5′- GTTCTGACTCCGCAGGAAATAGCGATGGTGCACCTAAGAACATGCAAG -3′ |
| $EA_{GsLL}$ Rev | 5′- GAGGGTCGGCCACCGACCCGTTTCGATCCCCATCGCTTGGAG -3′ |
